# Gemigliptin alleviates succinate induced endoplasmic reticulum stress and activation of hepatic stellate cells

**DOI:** 10.1101/2022.12.01.518797

**Authors:** Dinh-Vinh Do, Giang Nguyen, So Young Park, Eun-Hee Cho

**Affiliations:** Department of Internal Medicine, School of Medicine, Kangwon National University, Republic of Korea

**Keywords:** Succinate, Hepatic Stellate Cells, Endoplasmic Reticulum Stress, Gemigliptin, Liver Fibrosis

## Abstract

**Background:** Hepatic stellate cells (HSCs) activation is the principal event in the development of liver fibrosis in which succinate-GPR91 signaling has recently been shown to be a contributor. Moreover, endoplasmic reticulum (ER) stress has been reported to involve in HSC activation, but its association with succinate in pathogenesis of liver fibrosis remains scarce. In this study, we investigated the role of gemigliptin, an antidiabetic DDP-4 inhibitor, in the succinate-induced ER stress and activation of HSCs.

**Methods:** LX-2 cells, the immortalized human HSCs, were treated with succinate and gemigliptin. For animal experiments, C57BL/6N mice were divided into 3 groups: control diet, high-fat high-cholesterol (HFHC) diet, and HFHC diet mixed with gemigliptin.

**Results:** Succinate significantly induced HSC activation and increased expression of inflammatory markers and the increase in the migration of HSCs. The treatment of succinate also caused ER dilation and activated the unfolded protein response (UPR) signaling as PERK, eIF2alpha, Bip, suggesting increasing ER stress in HSCs. All responses of HSCs to succinate were attenuated with the co-treatment of gemigliptin. Moreover, the exposure of HSCs to tunicamycin, an inducer of ER stress, promoted the expression of α-SMA, proliferation and migration of HSCs. In vivo, the level of fibrotic and ER stress markers was increased in mice fed with HFHC diet and the administration of gemigliptin improved these changes in HFHC-induced mice.

**Conclusion:** This study showed the involvement of ER stress in the activation of succinate-induced LX-2 HSCs and gemigliptin significantly reduced ER stress in HSC activation. Therefore, gemigliptin may become an anti-fibrotic agent and targeting to succinate and ER stress may be a promising therapeutic in the management of liver fibrosis.

## INTRODUCTION

Chronic liver diseases can progress to liver fibrosis as an excessive wound-healing response to liver injury. In pathogenesis of liver fibrosis, hepatic stellate cells (HSCs) are mostly responsible for the accumulation of extracellular matrix (ECM) which leads to the formation of scar tissue. During injury, quiescent HSCs transdifferentiate into activated myofibroblasts, which is characterized by proliferation, chemotaxis, fibrogenesis, contractility, chemoattractant and cytokine release [1,2]. Numerous studies have depicted the underlying mechanism of HSC activation, senescence and the reversion, apoptosis of activated HSCs [2,3]. Succinate, an intermediate formed in the TCA cycle, has been demonstrated to involve in HSC activation in liver fibrosis [4–6]. Succinate accumulation via SIRT3-succinate dehydrogenase (SDH) pathway stimulates HSC activation [7,8]. Furthermore, succinate receptor, known as G-protein coupled receptor 91 (GPR91), was up-regulated in activated HSCs. The inhibition of the GPR91 expression via small interfering RNA and GPR91 antagonist was shown to deactivate HSCs [4,9]. These suggest the succinate-GPR91 signaling pathway has important role in regulation liver fibrosis.

The accumulation of unfolded or misfolded proteins in endoplasmic reticulum (ER) lumen leads to the activation of downstream signaling pathway of ER stress, called the unfolded protein response (UPR), to maintain the cellular homeostasis. It is known that ER stress is associated with the pathogenesis of liver diseases and hepatic fibrosis through the activation of UPR in multiple cell types, predominantly hepatocytes and HSCs [10,11]. Upon HSC activation in response to potent inducer, transforming growth factor β (TGFβ), ER stress and UPR signaling was noted in these studies [12–14]. This suggests UPR induction may be a result from the increase in production and secretion of ECM components in HSCs. On the other hand, UPR activation also causes fibrogenesis in HSCs and therefore ER stress may be a driver of HSC activation [11]. These observations show that ER stress plays a crucial role in HSC activation as well as liver fibrosis.

DPP-4 inhibitors were developed and approved as the therapeutic agents for treatment of type 2 diabetes [15]. Besides the role in diabetes treatment, DPP-4 inhibitors have been studied for their nonglycemic benefits on other diseases and conditions such as inflammation, cardiovascular and renal diseases [16–19]. Previous studies have shown that sitagliptin, linagliptin, vildagliptin, gemigliptin, and alogliptin could have beneficial effects on hepatic steatosis in observational human and/or animal-model data [20–26]. Moreover, DPP-4 inhibition with linagliptin, anagliptin attenuated Western diet-induced liver fibrosis in rodents [27–29]. The protective effects of DPP-4 inhibitors on liver fibrosis could be independent of glucose reduction, as evidenced by the administration of DPP-4 inhibitors in carbon tetrachloride rodent models [30–32]. To be more specific, sitagliptin and alogliptin were reported to suppress HSC activation process via Smad2/3 pathway [33,34], suggesting the involvement of DPP-4 inhibitors in protective effects on HSCs. In relation to ER stress, gemigliptin was shown to reduced tunicamycin-induced ER stress in H9c2 cardiomyocytes [35]. Taken together, we investigated the link between succinate and ER stress in HSCs and the role of gemigliptin in the succinate-induced ER stress and activation of HSCs.

## MATERIALS AND METHODS

### 1. Materials

The reagents used in this study were obtained from the indicated suppliers: succinate, tunicamycin, and antibodies against collagen type I (Col-I) from Sigma (St. Louis, MO, USA); antibodies against TGF-β, IL-6, p-PERK, p-eIF2α, IRE1α, CHOP, Bip, and PDI from Cell Signaling Technology (Richmond, CA, USA); antibodies against GAPDH from GeneTex (Irvine, CA, USA); antibodies against GPR91 from (Santa Cruz, CA, USA); antibodies against α-SMA and HIF-1α from Abcam (Cambridge, England); Transwell filters from Costar (Corning, NY, USA). All other materials were obtained from Sigma (St. Louis, MO, USA).

### 2. Cell culture

Human hepatic stellate cell line, LX-2 cells were kindly provided by Professor Ja June Jang, Seoul National University. Cells were maintained in Dulbecco’s Modified Eagle’s Medium (DMEM) containing 100 ml/L of fetal bovine serum (FBS) with 100,000 U/L of penicillin and 100 mg/L of streptomycin.

### 3. Western blot analysis

Cells were lysed in RIPA buffer (ATTO corporation, Tokyo, Japan). The protein contents of total cell lysates were determined using a BCA protein assay kit (Thermo Scientific Pierce, Rockford, IL, USA). The equal amounts of protein were separated by sodium dodecyl sulfate polyacrylamide gel electrophoresis (SDS-PAGE), and then transferred to a polyvinylidene fluoride (PVDF) membrane (Millipore, Bedford, MA, USA). After blocking with 5% non-fat dry milk in Tirs buffered saline containing 0.1% Tween-20 for 1 h at room temperature, the membranes were incubated with primary antibodies according to the instructions overnight at 4°C or 2 h at room temperature. The membranes were then incubated with the appropriate horseradish peroxide conjugated secondary antibodies at room temperature. Signals were detected via enhanced chemiluminescence, using Westsave Star Detection Reagent system (AbFrontier, Seoul, Korea). The band intensity was calculated using the Image J program [Ver 1.42, National Institute of Health (NIH) Image, developed and maintained by the NIH (Bethesda, MD, USA)].

### 4. Real-time PCR

Total RNA was isolated using RNeasy plus micro kit (QIAGEN, Hilden, Germany) or TRIzol reagent (Roche, Indianapolis, IN, USA). After extraction of total RNA, cDNA was synthesized from 1 μg of total RNA with a Maxime RT PreMix kit (iNtRON Biotechnology, Gyeonggi, Korea) according to the manufacturer’s instructions. Real-time PCR analysis was performed with SYBR Green using QuantTudio™6 Flex Real-Time PCR System. The relative mRNA levels of target genes were assessed by using the 2^_ΔΔCt^ method.

### 5. Cell viability assay

LX-2 cells were plated in 24-well plates with DMEM containing 100 mL/L FBS at 37°C. After 24 h, the medium was replaced with several concentration gemigliptin in the absence or presence of 1600 μmol/L of succinate in fresh DMEM containing 100 ml/L of FBS. Viable cell numbers were estimated via CCK-8 (Dojindo Molecular Technologies, Inc., Rockvile, USA), in accordance with the manufacturer’s instructions. Succinate was dissolved in phosphate buffered saline (PBS).

### 6. Cell migration assay

For the wound migration assay, LX-2 cells were plated at 5 × 10^5^ cells/well in DMEM containing 100 ml/L of FBS into 6well plate. Cells at 90% confluence were incubated for 1 h with 1 mg/L of mitomycin C. After that, injury line was made using a yellow tip, and the cell monolayers were washed gently with PBS. The cells were then incubated for 6 h with several concentration gemigliptin in the absence or presence of 1600 μmol/L of succinate in fresh DMEM containing 100 ml/L of FBS. Cell migration was measured with microscopy at 0 and 6h, and the measured widths of the injury lines were plotted as moving distance.

For the transwell migration assays, LX-2 cells (3 × 10^4^ cells/filter) were then plated onto transwell filters (Corning, NY, USA) in a 24-well plate. The transwell filter was precoated with 10 μg Type IV collagen. The lower chambers of the wells were filled with DMEM containing 15% FBS as a chemoattractant. The cells were incubated for 6 h with several concentration gemigliptin in the absence or presence of 1600 μmol/L of succinate in fresh DMEM containing 100 ml/L of FBS. Migrated cells were fixed and stained with hematoxylin and eosin. Migrated cell numbers in 8 separate fields were counted using light microscopy at 200× magnification.

### 7. TEM

LX-2 cells were plated and treated with several concentration gemigliptin in the absence or presence of 1600 μmol/L of succinate for 24h. 24h incubated cells were fixed with 2% glutaraldehyde and 2% paraformaldehyde in phosphate buffer (pH 7.4) for 1 h at 4°C and then postfixed osmium tetroxide for 40 min at 4°C. The cells were dehydrated in a graded series of ethanol. The cells were treated with graded propylene oxide series and embedded into Epon. The sections were then ultra-thin sectioned in 80-nm and placed on copper grid. The final samples were stained with uranyl acetate and lead citrate. After that the samples were observed using a transmission electron microscope (JEOL-2100F, USA, 200kV) at the Korea Basic Science Institute, Chuncheon.

### 8. Animal experiments

This animal study was approved by the Institutional Animal Care and Use Committee of Kangwon National University and conducted in accordance with its guidelines (Protocol approval # KW-180103-1). Three-week old, male C57BL/6N mice were purchased from Doo Yeol Biotech (Seoul, Korea). All mice were housed at ambient temperature (22 ± 1 °C) with a 12/12-h light/dark cycle and with free access to water and food in The National Kangwon University animal care facility. The mice were randomly divided into 3 groups: 1) the Control group was fed with control diet (2018S, ENVIGO, USA) 2) the NASH model group fed with HFHC diet, and 3) the treatment group was fed with HFHC diet mixed with 0.4% gemigliptin. The nutritional compositions of the HFHC are reported in Supplementary Table 1. The study duration was 8 weeks. The blood was collected, and plasma was isolated by centrifugation. All tissue samples were stored at −80°C.

For histological analysis samples of mouse liver were fixed in 4% paraformaldehyde for IHC analysis. After that dehydration through a graded series of ethanol solutions, the tissues were embedded in paraffin wax. Serial frontal sections were cut and stained with hematoxylin and eosin (H&E), and Masson’s trichrome. Photographs were obtained using a microscope.

### 9. Statistical analyses

All data are expressed as mean ± SEM. Data analyzed via analysis of variance. Differences between the treatment groups were evaluated via Duncan’s multiple range test, using SAS for Windows version 9.2 software (SAS Institute, Cary. NC, USA). Differences were considered significant at P < 0.05.

## RESULTS

### 1. Gemigliptin inhibited activation and GPR91 expression of succinate-induced HSCs

The expression of fibrotic markers, collagen type I and α-SMA, succinate receptor – GPR91 and inflammatory markers were significantly increased by the treatment of succinate 1,600 μmol/L for 24 h in LX-2 cells. Particularly, gemigliptin suppressed all these changes in succinate-induced LX-2 cells in dose-dependent manner (Fig. 1.).

**Figure 1.**
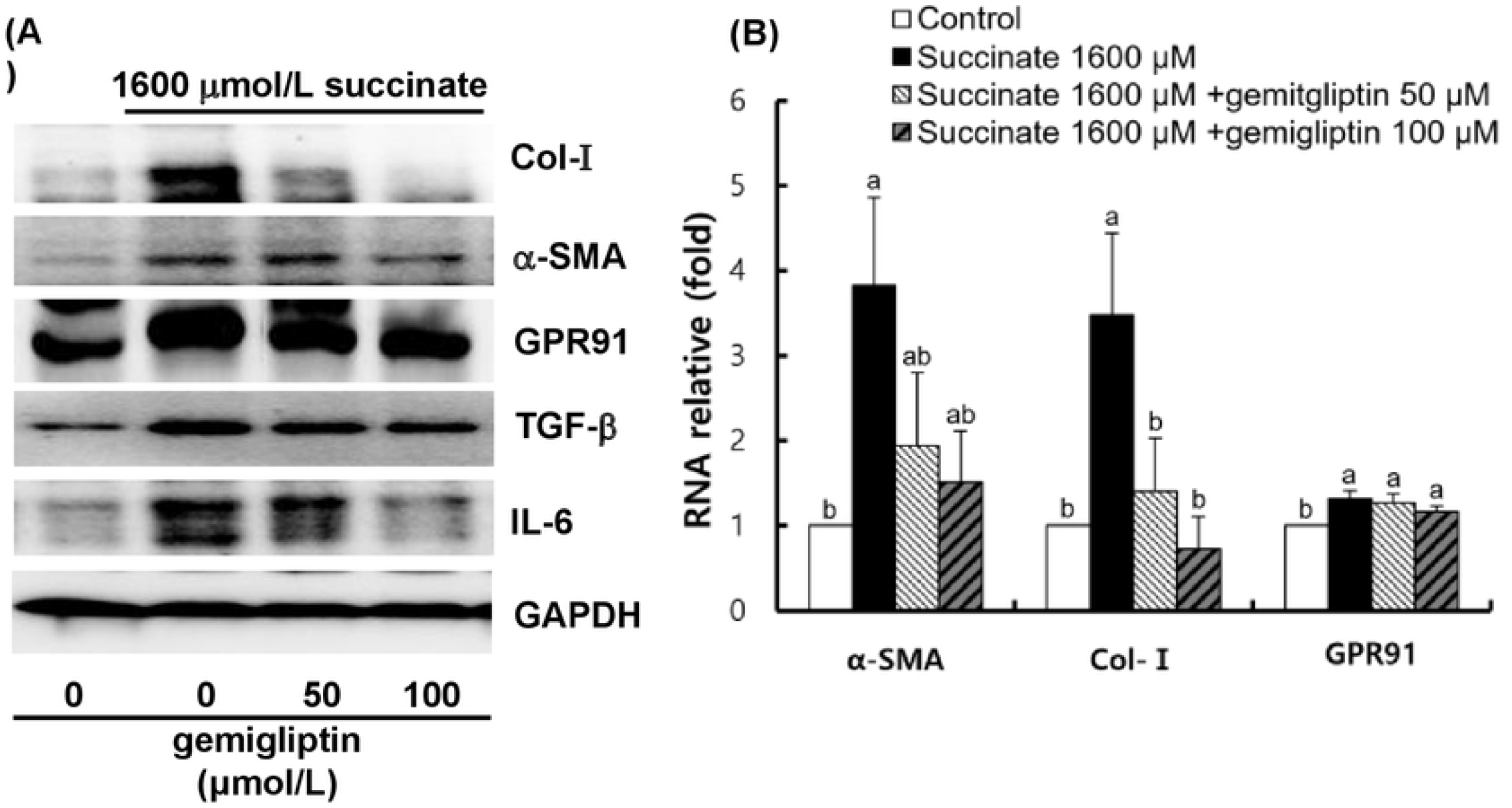
Gemigliptin inhibited activation and GPR91 expression of succinate-induced HSCs. LX-2 cells were treated with 0–100 μmol/L gemigliptin with or without succinate 1600 μmol/L for 24 h. **(A)**. Representative images from Western blotting analysis of the protein expressions of collagen type I, α-SMA, GPR91 and GAPDH in LX-2 cells after 24 h treatment from three independent experiments. **(B)**. Real-time PCR analysis of relative mRNA expression of α-SMA, collagen type I and GPR91 normalized to GAPDH in LX-2 cells after 24 h treatment. Each bar represents the mean ± SEM (n = 3). Means with different letters differ significantly, *P* < 0 .05.

### 2. Gemigliptin inhibited succinate-stimulated HSC migration

As expected, succinate treatment significantly increased wound healing migration (Fig. 2A.) and Transwell migration (Fig. 2B.) in LX-2. Gemigliptin co-treatment decreased level of HSC migration in LX-2 cells, which had stronger effect at 100 μmol/L gemigliptin.

**Figure 2.**
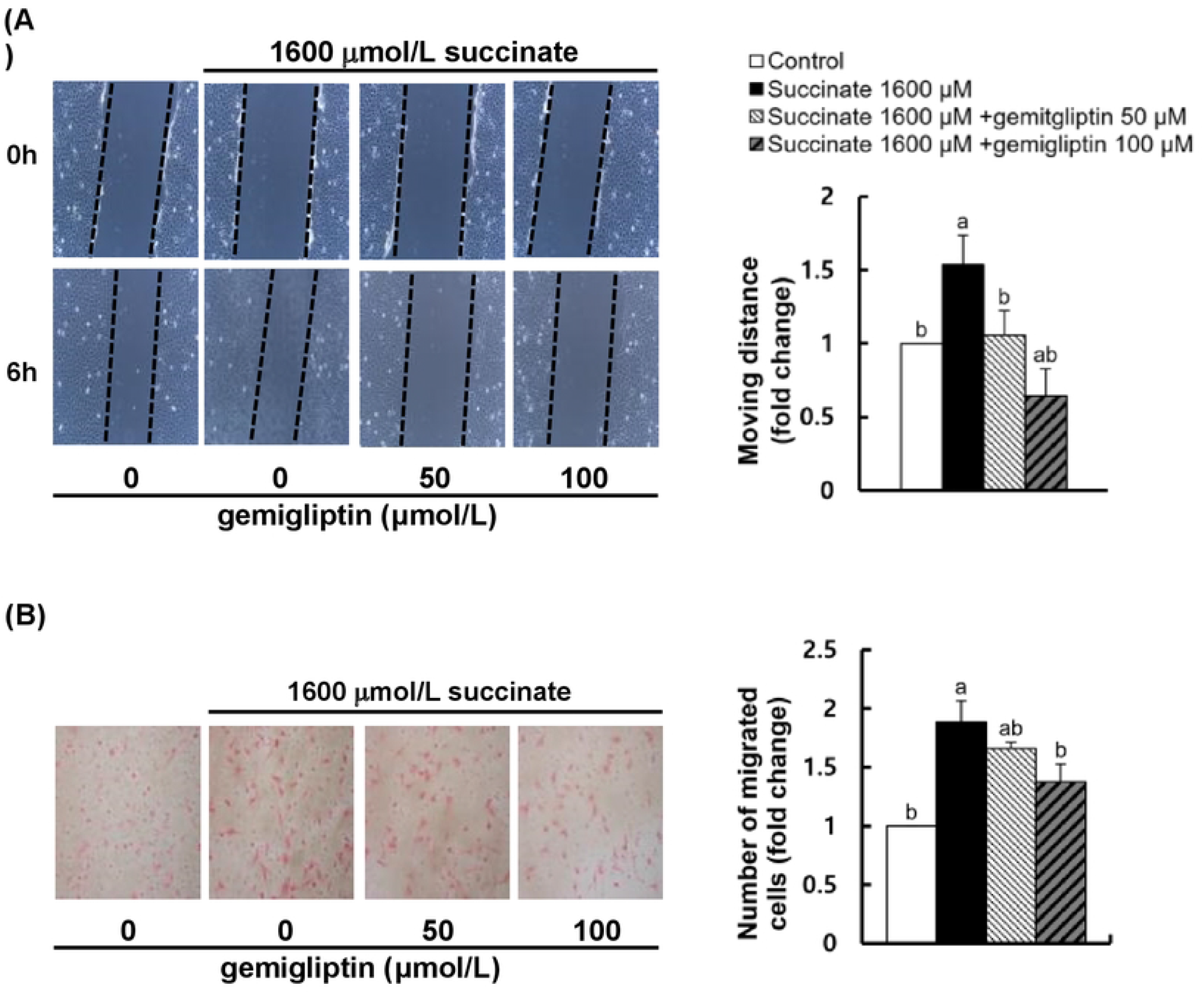
Gemigliptin inhibited succinate-stimulated HSC migration. **(A-B)** LX-2 cells were treated with 0–100 μmol/L gemigliptin with or without succinate 1600 μmol/L for 6 h to evaluate cell migration. **(A)** Representative images of wound healing migration and statistical analysis shown as relative moving distance from three independent experiments. (**B**) Representative images of Transwell migration and statistical analysis shown as relative migrated cells from three independent experiments. Each bar represents the mean ± SEM (n = 3). Means with different letters differ significantly, *P* < 0 .05.

### 3. Gemigliptin inhibited succinate-stimulated ER stress in HSCs

We investigated whether succinate could cause ER stress in HSCs by evaluating ER structure of LX-2 cells and the expression of ER stress-related markers. As shown in Fig. 3A., the swollen ER was markedly noted in succinate-treated LX-2 cells, which was reversed by gemigliptin 100 μmol/L treatment. The expression of UPR signaling (p-PERK, p-eIF2α, IRE1α, CHOP, Bip except PDI) was also upregulated by succinate treatment but decreased when co-treatment with gemigliptin (Fig. 3B-C.). These results suggested that succinate may induce ER stress which was inhibited by gemigliptin. In addition, we observed that HIF-1α, recognized as pro-fibrotic factor and stabilized and increased by succinate treatment and gemigliptin reduced the expression of HIF-1α.

**Figure 3.**
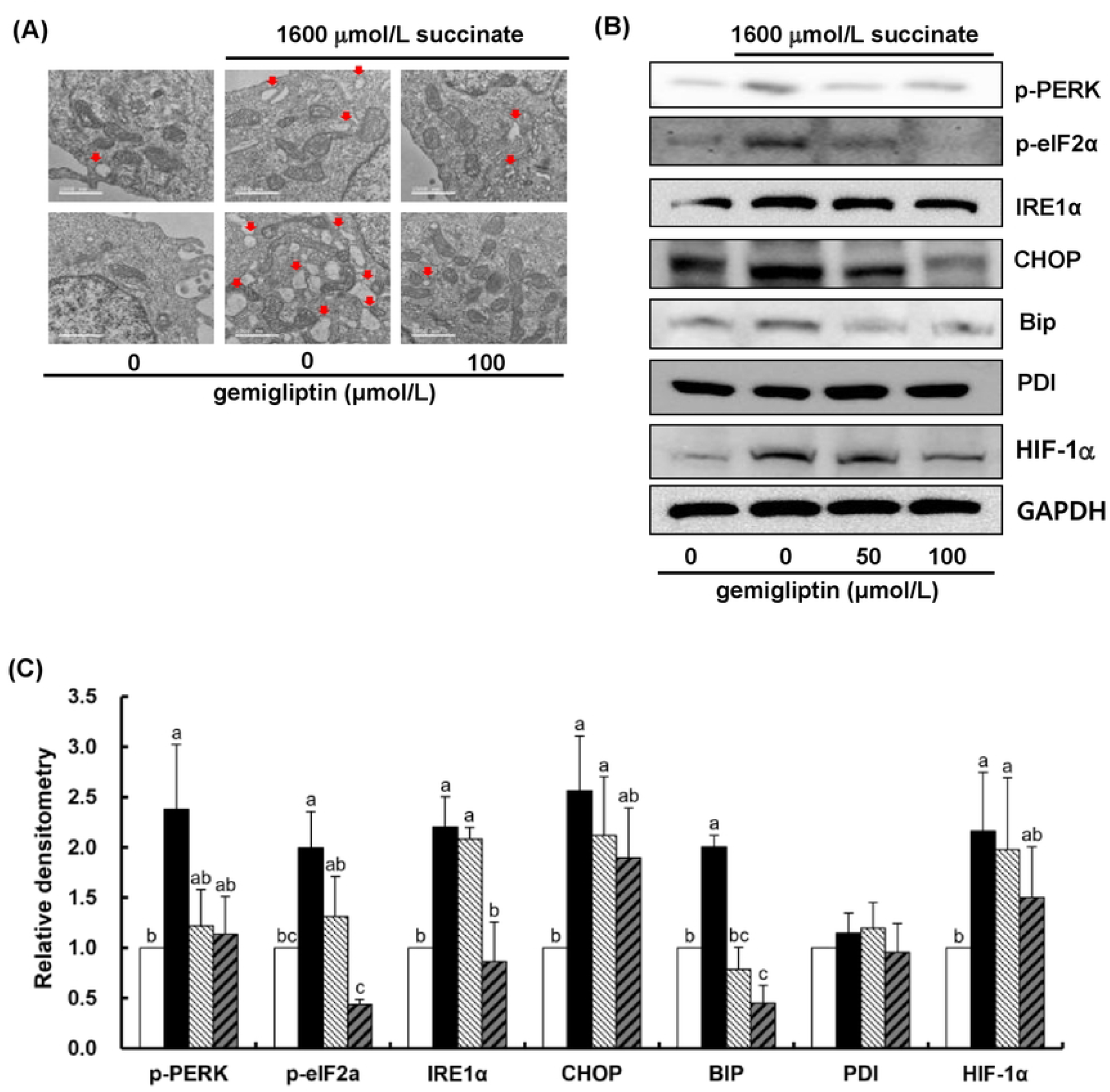
Gemigliptin inhibited succinate-stimulated ER stress in HSCs. **(A-B)** LX-2 cells were treated with 0–100 μmol/L gemigliptin with or without succinate 1600 μmol/L for 24 h. **(A)** Electron microscopic images showing the ultrastructure of LX-2 cell; solid arrows highlight the swollen endoplasmic reticulum. **(B-C)**. Representative images (**B**) and relative densitometric analysis (**C**) from Western blotting analysis of the protein expressions of p-PERK, p-eIF2α, IRE1α, CHOP, Bip, PDI, HIF-1α and GAPDH in LX-2 cells after 24 h treatment from three independent experiments with protein expression normalized to GAPDH. Each bar represents the mean ± SEM (n = 3). Means with different letters differ significantly, *P* <0 .05.

**Figure 4.**
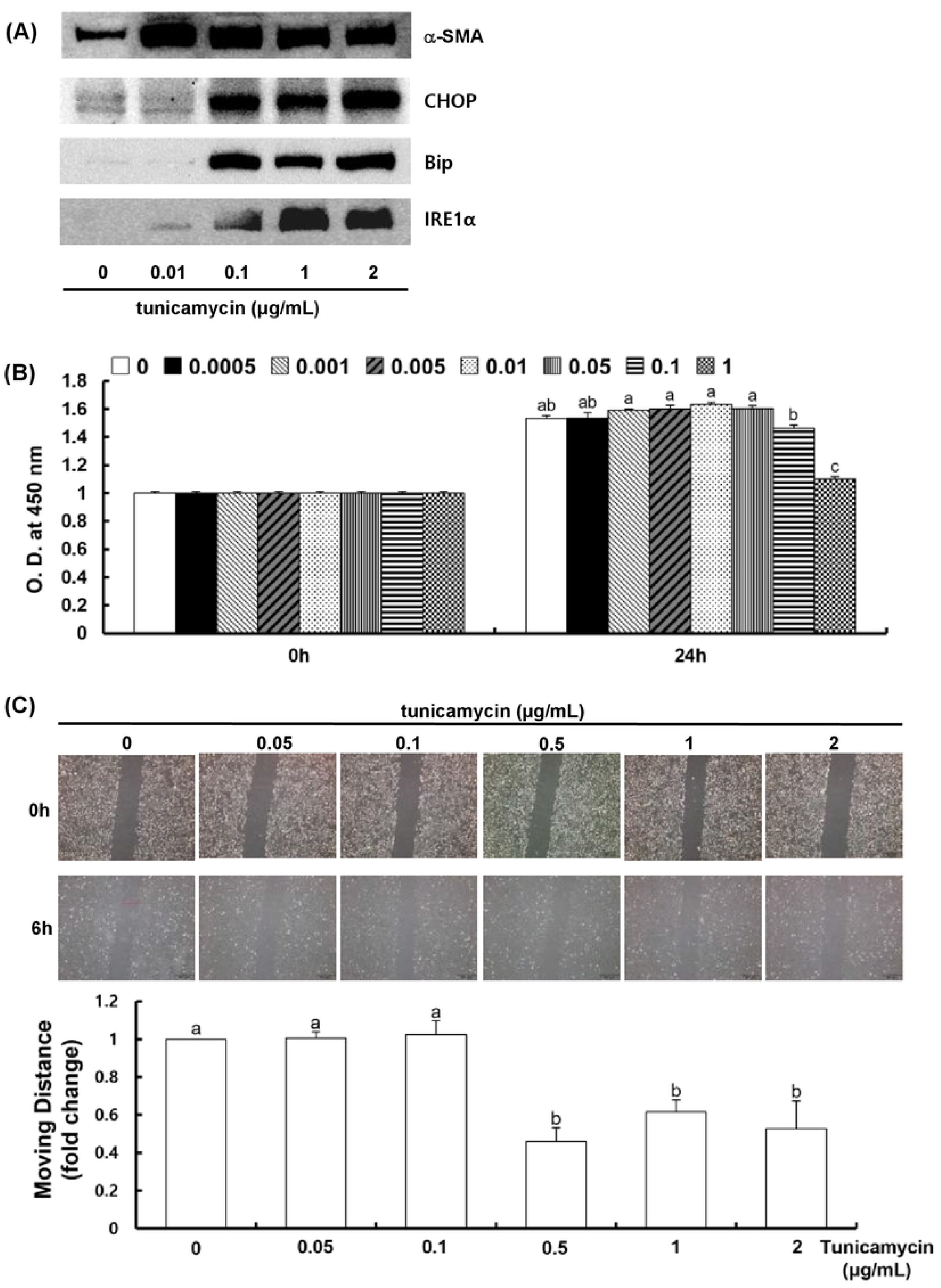
Tunicamycin at low concentration stimulated the expression of α-SMA and ER stress-related proteins in HSCs. LX-2 cells were treated with tunicamycin 0-2 μg/mL for 24 h. **(A)** Representative images from Western blotting analysis of the protein expressions of α-SMA, CHOP, Bip, IRE1α and GAPDH in LX-2 cells after 24 h treatment from three independent experiments. **(B)** The viability of LX-2 cells shown as optical density at 0 h and 24 h treatment with tunicamycin. **(C)** Representative images of wound healing migration and statistical analysis shown as relative moving distance from three independent experiments. Each bar represents the mean ± SEM (n = 3). Means with different letters differ significantly, *P* <0 .05.

### 4. Tunicamycin at low concentration stimulated the expression of α-SMA and ER stress-related proteins in HSCs

To further investigate the role of succinate in ER stress and HSC activation, we treated LX-2 cells with strong ER-stress inducer, tunicamycin, with several concentrations. ER stress was observed in LX-2 cells after tunicamycin treatment with low and high concentration by the up-regulation of CHOP, Bip, IRE1α. The results also showed that tunicamycin induced HSC activation as evidenced by the increase in α-SMA expression and proliferation of LX-2 cells. However, we noted a decrease in viability of LX-2 cells with tunicamycin treatment at concentration greater than 0.1 μg/mL. The migration of LX-2 cells also decreased with tunicamycin greater than 0.5 μg/mL.

### 5. Gemigliptin improved fibrotic liver and ER stress in HFHC diet-induced mice

We evaluated the effect of gemigliptin on liver fibrosis in HFHC diet-induced mouse model. After 8-week administration of HFHC diet, we observed hepatic steatosis and fibrosis in mice as shown by H&E and Masson’s-trichrome staining (Fig. 5A.). The expression of collagen type I and α-SMA was increased in HFHC group which was suppressed by gemigliptin addition. Moreover, HFHC diet upregulated the expression of ER stress markers, including CHOP, Bip, p-eIF2α, which was also inhibited by gemigliptin.

**Figure 5.**
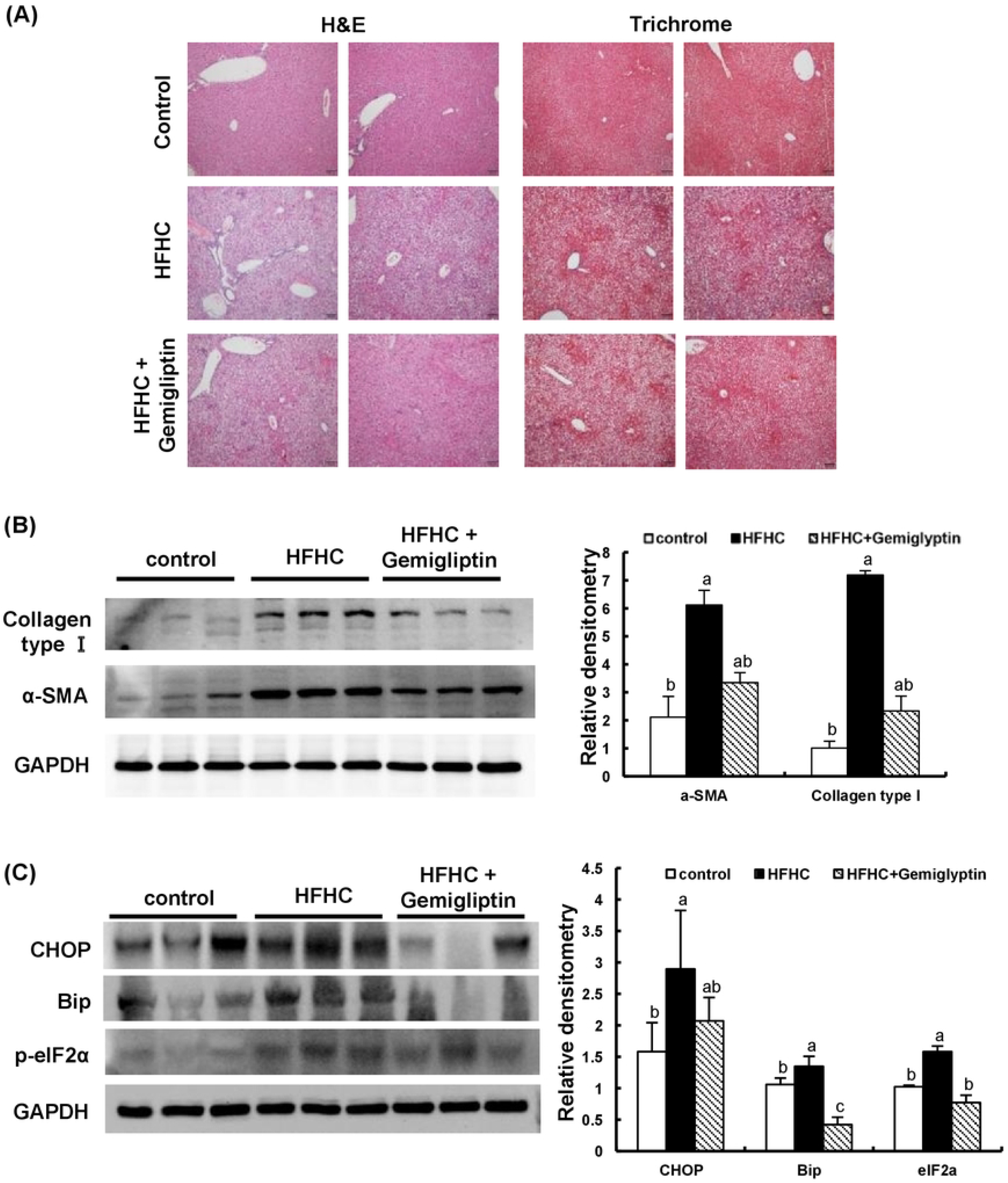
Gemigliptin improved fibrotic liver and ER stress in HFHC diet-induced mice. Mice were fed with control diet, HFHC diet or HFHC diet with gemigliptin for 8 weeks. **(A)** Representative Hematoxylin & Eosin staining and Masson’s trichrome staining of mouse livers. All slides are magnified 400X. **(B-C)** Representative images and relative densitometric analysis from Western blotting analysis of the protein expressions of collagen type I, α-SMA, GAPDH **(B)** and CHOP, Bip, p-eIF2α, GAPDH **(C)** in mouse liver samples with protein expression normalized to GAPDH. Each bar represents the mean ± SEM (control n=5, others n=10). Means with different letters differ significantly, *P* <0 .05.

## DISCUSSION

In this study, we evaluated the effects of succinate and DPP-4 inhibitor, gemigliptin, in relation to ER stress and activation of HSCs and livers of mice fed with HFHC diet. We found that HSC activation and ER stress were induced by succinate and ER stress induction could drive HSCs to activated phenotype. These changes in HSCs caused by succinate were effectively ameliorated by the treatment of gemigliptin. The beneficial effects of gemigliptin on liver fibrosis and ER stress were also noted in HFHC diet-induced mice.

HSC activation is one of the key events during progression of liver fibrosis, which can be induced by several fibrogenic mediators. Succinate and succinate-GPR91 receptor signaling have been found to contribute to HSC activation in the close relation to hepatocytes. Studies showed that succinate was released into extracellular space resulting from succinate accumulation in hepatocytes. Succinate can act as a signaling molecule which activate GPR91 receptor and subsequently induce HSC activation [4–7]. Our results have confirmed the effects of succinate on HSCs that succinate induced up-regulation of fibrogenic, inflammatory markers and migration in LX-2 cells. Remarkably, we observed that a DPP-4 inhibitor, gemigliptin, reversed these changes in LX-2 cells in a dose-dependent manner and reduced liver fibrosis in mice fed with HFHC diet. Several previous studies also reported that other DPP-4 inhibitors had beneficial role in liver fibrosis, especially sitagliptin and alogliptin could inhibit HSC activation which was induced by potent fibrogenic inducers such as transforming growth factor (TGF)-β1 and platelet-derived growth factor [33,34]. In our study, we noted that protein expression of GPR91 was up-regulated by succinate treatment and gemigliptin attenuated this effect, suggesting that succinate-GPR91 signaling pathway may be involved in gemigliptin action on HSCs.

Extensive evidence has shown the close relationship between ER stress and liver fibrosis, in which chemical-induced ER stress could activate HSCs and alternatively HSC activation could regulate ER stress through UPR signaling [11]. The activation of UPR is mediated through three ER stress sensors, including inositol requiring enzyme 1 α (IRE1α), protein kinase RNA-like ER kinase (PERK), and activating transcription factor 6 α (ATF6α) [10,11]. Besides that, some DPP-4 inhibitors were reported to suppress ER stress in different organs of several models. In rodent livers, sitagliptin, saxagliptin and linagliptin were shown to attenuated markers of ER stress which was induced by methionine/choline-deficient diet, fructose and high fat diet, respectively [23,36,37]. Particularly, vildagliptin reduced ER dilation and UPR signaling in the liver of high fat diet mice [38]. In HepG2 cells, ER stress induced by palmitic acid was also attenuated by the treatment of vildagliptin or VLATSGPG (a peptide known to inhibit dipeptidyl peptidase IV) [38,39]. In our study, we observed that in LX-2 HSCs, succinate treatment caused ER dilation and the result was further confirmed by the changes in UPR signaling, including the increase in IRE1α, phosphorylation of PERK-eIF2α, CHOP, Bip, showing link between succinate-GPR91 pathway and ER stress. Additionally, we noted that the co-treatment with gemigliptin decreased the state of ER stress concurrently with activation of LX-2 cells. These results are consistent with previous studies on ER stress amelioration of DPP-4 inhibitors [23,36–38].

In addition, tunicamycin treatment in our study induced activation of LX-2 cells at low dose of 0.01 μg/mL though there was no significant difference in IRE1α, Bip and CHOP expression. At high doses, tunicamycin increased ER stress markers but it reduced viability and migration of LX-2 cells. Tunicamycin is known as a potent ER stress inducer which can lead to HSC activation [40] but severe UPR with high dosage can also result in apoptosis [41,42]. One study showed that primary mouse HSCs which were cultured on plastic dish underwent activation from day fourth while UPR genes were induced just after 2 h. UPR was also seen earlier than the up-regulation of activation markers in freshly isolated HSCs from mice received single dose of CCl_4_, suggesting UPR is expressed early during HSC activation [43]. Therefore, we speculated that succinate in our study may act like an ER stress inducer like low-dose tunicamycin, which subsequently activate HSCs. However, this implication as well as the expression of UPR and HSC activation at different time point with succinate require more confirmation.

The data on relationship between succinate-GPR91 and ER stress in liver diseases remains sparse. A previous study reported that in ER-stressed mice induced with tunicamycin, GPR91 was one of the differentially expressed genes that predominantly identified from liver tissues [44]. In our study, beside the changes in ER stress, succinate treatment in LX-2 cells also increased the expression of HIF-1α, a protein recognized as an important contributor to development of liver fibrosis [45]. In cytosol, HIF-1α is stabilized by succinate accumulation, leading to “pseudohypoxia” condition [46]. A previous study showed that extracellular succinate may be transported into cytosol and subsequently stabilizing and activating HIF-1α [47]. Moreover, growing literature has implicated the crosstalk between UPR and hypoxia in which HIF-1α is involved in regulation of ATF4 and ATF6α/XBP1s, and HIF-1α stabilization may promote UPR signaling [48,49]. Therefore, the involvement of HIF-1α signaling may be one of mechanisms to explain how succinate treatment in our study induced ER stress and activation in HSCs, but further studies are required to confirm and give in-depth mechanism.

In conclusion, we showed that succinate treatment induced ER stress and activation in LX-2 cells and gemigliptin significantly reversed these changes. Therefore, succinate and ER stress continue to be the target of liver fibrosis resolution and gemigliptin may become an anti-fibrotic agent.

---

**Supplementary table 1.** The composition of experimental diet.

| Composition | Control | HFHC | HFHC+Gemigliptin |
| --- | --- | --- | --- |
| 2018S (g) | 100 | 38.25 | 38.25 |
| Cocoa butter (g) | — | 60 | 60 |
| Cholesterol (g) | — | 1.25 | 1.25 |
| Cholate (g) | — | 0.5 | 0.5 |
| Gemigliptin (g) |  |  | 0.4 |
| Total (g) | 100 | 100 | 100.4 |

